# Spatial contextual information modulates affordance processing and early electrophysiological markers of scene perception

**DOI:** 10.1101/2024.03.15.585247

**Authors:** Clément Naveilhan, Maud Saulay-Carret, Raphaël Zory, Stephen Ramanoël

## Abstract

Scene perception allows humans to extract information from their environment and plan navigation efficiently. The automatic extraction of potential paths in a scene, also referred to as navigational affordances is supported by scene-selective regions (SSRs) that enable efficient human navigation. Recent evidence suggests that the activity of these SSRs can be influenced by information from adjacent spatial memory areas. However, it remains unexplored how these contextual information could influence the extraction of bottom-up information, such as navigational affordances, from a scene and the underlying neural dynamics. Therefore, we analyzed event-related potentials (ERPs) in 26 young adults performing scene and spatial memory tasks in artificially generated rooms with varying numbers and locations of available doorways. We found that increasing the number of navigational affordances only impaired performance in the spatial memory task. ERP results showed a similar pattern of activity for both tasks, but with increased P2 amplitude in the spatial memory task compared to the scene memory. Finally, we reported no modulation of the P2 component by the number of affordances in either task. This modulation of early markers of visual processing suggests that the dynamics of SSR activity are influenced by a priori knowledge, with increased amplitude when participants have more contextual information about the perceived scene. Overall, our results suggest that prior spatial knowledge about the scene, such as the location of a goal, modulates early cortical activity associated with scene-selective regions, and that this information may interact with bottom-up processing of scene content, such as navigational affordances.

## Introduction

Spatial navigation is a fundamental cognitive ability that allows us to find our way and move through complex and diverse environments. This high-level cognitive function relies on the integration of information from multiple sensory modalities, among which vision, and particularly the processing of a visual scene, plays a central role for humans (Dilks et al., 2022; Ekstrom, 2015; Epstein & Baker, 2019; Julian et al., 2018). Through visual perception, we are able to rapidly integrate a large quantity of spatially relevant information from a scene, ranging from low- and mid-level properties (*i*.*e*., edges, orientations, spatial layout, etc.) to high-level properties (*i*.*e*., conceptual attributes, affordances), which allow us to navigate efficiently in our environment (Cichy et al., 2017; Greene & Hansen, 2020; Harel et al., 2022; Kauffmann et al., 2015)

This interplay between scene perception and navigation is subtended by the involvement of three scene-selective brain regions (SSRs). The parahippocampal place area (PPA) in the parahippocampal cortex plays a role in scene categorization, retrieval of navigationally relevant cues, and sensitivity to spatial layout (Epstein, 2008; Janzen & van Turennout, 2004; Persichetti & Dilks, 2018; Sun et al., 2021). The retrosplenial complex (RSC), also referred to as the medial place area (Silson et al., 2016), in the posterior cingulate cortex, is associated with map-based navigation, and involves the computation of heading directions and the translation of information between egocentric and allocentric spatial reference frames (Alexander et al., 2023; Mitchell et al., 2018; Vann et al., 2009). Finally, the occipital place area (OPA) near the transverse occipital sulcus has been reported to support first-person vision-guided navigation through the encoding of egocentric distance and local scene elements (Dilks et al., 2013; Julian et al., 2016; Kamps et al., 2016; Persichetti & Dilks, 2018).

Among these three SSRs, the OPA region stands out as particularly interesting when considering the relationships between scene processing and navigation in a surrounding environment. Indeed, this region is highly sensitive to sensory inputs associated with egocentric distance (Persichetti & Dilks, 2016) and to first-person egocentric motion cues (Kamps et al., 2016; Pitcher et al., 2019). The OPA also plays a specific role in encoding stimuli that constrain potential movement through the environment, including boundaries, isolated obstacles, and navigational affordances (Bonner & Epstein, 2017; Dillon et al., 2018; Harel et al., 2022; Henriksson et al., 2019; Kamps et al., 2016; Park & Park, 2020). Importantly, it has been proposed that the OPA could automatically extracts navigational affordances from the passive perception of a visual scene, even when participants are not engaged in a navigation task (Bonner & Epstein, 2017). This proposed mechanism of encoding the action possibilities in a scene was further investigated by Harel *et al*. (2022) using electroencephalography (EEG). They analyzed event-related potentials (ERPs) and notably the P2 component recorded from occipito-parietal electrodes, which has been proposed to reflect the activity of SSRs during scene processing (Harel et al., 2016; Kaiser et al., 2020). The authors reported that the affordances available in a scene are already represented during the early stages of visual processing, with a modulation of the P2 component (*i*.*e*., occurring 200 ms after image onset) over occipito-parietal electrodes consistent with previous computational studies (Greene & Hansen, 2020; Groen et al., 2018). These findings from neuroimaging studies led to the proposal that the OPA region may support the automatic processing of navigational affordances through the local environment, integrating multiple navigation-related features and generating a “navigation file” (Kang & Park, 2023).

The notion that the OPA region is a bottom-up detector has also been expanded by Choi *et al*. (2020), who demonstrated the ability of this brain region to integrate both visual and semantic information in order to compute navigational possibilities in scenes beyond what can be inferred from visual features alone. Aminoff & Tarr (2021) provided further evidence supporting this point by reporting that the context in which a visual scene is perceived modulates the activity of the SSRs, reflecting the integration of both bottom-up and top-down information. The convergence of these two types of information has thus been proposed to facilitate efficient encoding of a scene (Kaiser & Cichy, 2021), with top-down predictions of an upcoming scene influencing even the earliest stages of its processing, and particularly the P2 component (McLean et al., 2023). In the same vein, a recent series of functional magnetic resonance imaging (fMRI) studies have shed light on the neural basis of this proposed interplay between bottom-up (*i*.*e*., more perceptual) and top-down (*i*.*e*., more mnemonic) information in SSRs (Steel et al., 2021, 2023, 2024).The authors identified three “place-memory areas” adjacent to and partially overlapping with the anterior part of the more perceptual PPA, RSC, and OPA regions, challenging conventional distinctions between perceptual and memory systems (Baldassano et al., 2016; Martin & Barense, 2023).

Despite converging evidence of both bottom-up and top-down integration across SSRs, the potential influence of prior visuospatial knowledge representations on cortical correlates of early visual processing remains unexplored (Bartnik & Groen, 2023). To address this caveat, the processing of navigational affordances offers an interesting experimental framework. Underpinned by an early automatic extraction by the OPA, the early processing of these affordances has been proposed to also be influenced by the participant’s action intentions when presented with a scene (Djebbara et al., 2019; Wang et al., 2023), in a similar way as the early perception of an object is influenced by semantic knowledge about it (Enge et al., 2023). However, it remains unclear how prior spatial knowledge about the scene might modulate the neural signature associated with the automatic perception of navigation affordances (Harel et al., 2022). In other words, in a navigation task involving multiple visible paths in a visual scene, how might prior knowledge about the location of a goal constrain the processing of available navigational affordances?

To address this question, we analyzed behavioral (*i*.*e*., reaction time and response accuracy) and event-related potentials (ERPs) recorded over occipito-parietal electrodes while participants performed scene memory and spatial memory tasks in static room images with a varied number and location of navigational affordances (hereafter also referred to as open doors). At the behavioral level, we hypothesize that a longer reaction time will be observed in the spatial memory task than in the scene memory task, reflecting the additional processing demands associated with spatial memory retrieval. At the cortical level, we hypothesize that during the spatial memory task, activity over electrodes related to SSR will be greater than in the scene memory task, associated with increased knowledge of the spatial context (*i*.*e*., knowledge of the path to a goal). Considering the processing of affordances, we expect to report a modulation of the P2 component by the number of navigational affordances during the scene memory task, similar to what was observed by Harel *et al*. (2022). However, during the spatial memory task, when participants have prior knowledge of the path to the goal, P2 activity is not expected to be modulated by the number of affordances available.

## Methods

### Participants

We recorded the EEG activity of 30 healthy young participants. All were right-handed, had normal or corrected-to-normal vision, and no history of neurological or cognitive disorders. The study was approved by the local ethics committee (CERNI-UCA opinion no. 2021-050) and participants gave their informed consent before participation. Four participants were excluded from the analysis due to excessive artifacts in their EEG recordings, as determined by signal-to-noise ratio calculations and visual inspection. The final sample size was 26 participants (mean age: 23.26 years; SD: 0.61; range: 19-31; 14 females). This sample size was determined by a priori power analysis using G*Power (version 3.1.9.7; Faul et al., 2007). The power analysis was based on data from a previous ERP study investigating behavioral and neural differences between young and older adults during a landmark-based reorientation task contrasted with a passive scene perception (Naveilhan et al., 2023). Analysis of the young participants’ data indicated that a minimum of 23 participants was required to achieve a power of 0.95 at an alpha level of 0.05.

### Stimuli and procedure

Visual stimuli were generated using Unity Engine (version 2019.2.0.0f1) and displayed on an iiyama ProLiteB2791HSU screen (resolution 1920x1080, refresh rate 30-83 Hz) positioned 60 cm from the participants. PsychoPy (version 2022.13) software running on a Dell Precision 7560 computer (equipped with an Intel® Xeon® W-11955 processor) was used to present the stimuli. The visual stimuli consisted of simple rectangular rooms resembling those used in previous studies (Bonner & Epstein, 2017; Harel et al., 2022). Each room featured either a door or a gray rectangle on each of its three visible walls. Seven door configurations (left, right, center, left-center, left-right, right-center, left-right-center, doors open) and seven variations with distinct wall colors were created, resulting in a total of 49 unique stimuli. The experimental paradigm consisted of six similarly structured blocks, presented in a randomized order to each participant (**Figure 1**). The stimuli were presented in a pseudorandomized order within each block to maintain participant engagement. The paradigm comprised two main conditions involving a *scene memory* task and a *spatial memory* task. In the *scene memory* task, participants were asked to identify whether the scene presented matched the previous scene (1-back task) using wall color as the only discriminative cue. They were instructed to press a key if the scene matched the previous one. In this phase, 64 images were presented corresponding to the four door configurations in two distinct environments (*i*.*e*., two rooms with different wall colors), each repeated 8 times. In the *spatial memory* task, participants first completed a learning phase, in which they passively navigated through the rooms at a speed of 3 virtual meters per second and a turning speed of 40°/s. They were instructed to memorize the path leading to a hidden goal located in one of the adjacent rooms (left, center, or right). During this phase, participants were exposed to two distinctively colored rooms matching those used in the scene memory task. They were tasked with associating each wall color with a specific goal direction (*e*.*g*., in the blue room, the goal was located on the right, while in the red room, it was on the left). We ensured that the color of the wall was the only cue that guided participants through both the scene and spatial memory tasks to have the same perceptual load in both tasks. During the spatial memory task, participants were presented with the same images as during the scene memory task, but had more contextual information (*i*.*e*., the position of a goal in an adjacent room). They were instructed to use a USB keyboard to indicate the direction of the goal as quickly and accurately as possible, to ensure that they had acquired the knowledge of the goal position. Importantly, the spatial memory task was performed after the scene memory task within each block to guarantee that participants had no prior knowledge of the goal’s location during the scene memory phase. We ensured that the door leading to the goal was open during the spatial memory task.

**Figure 1.**
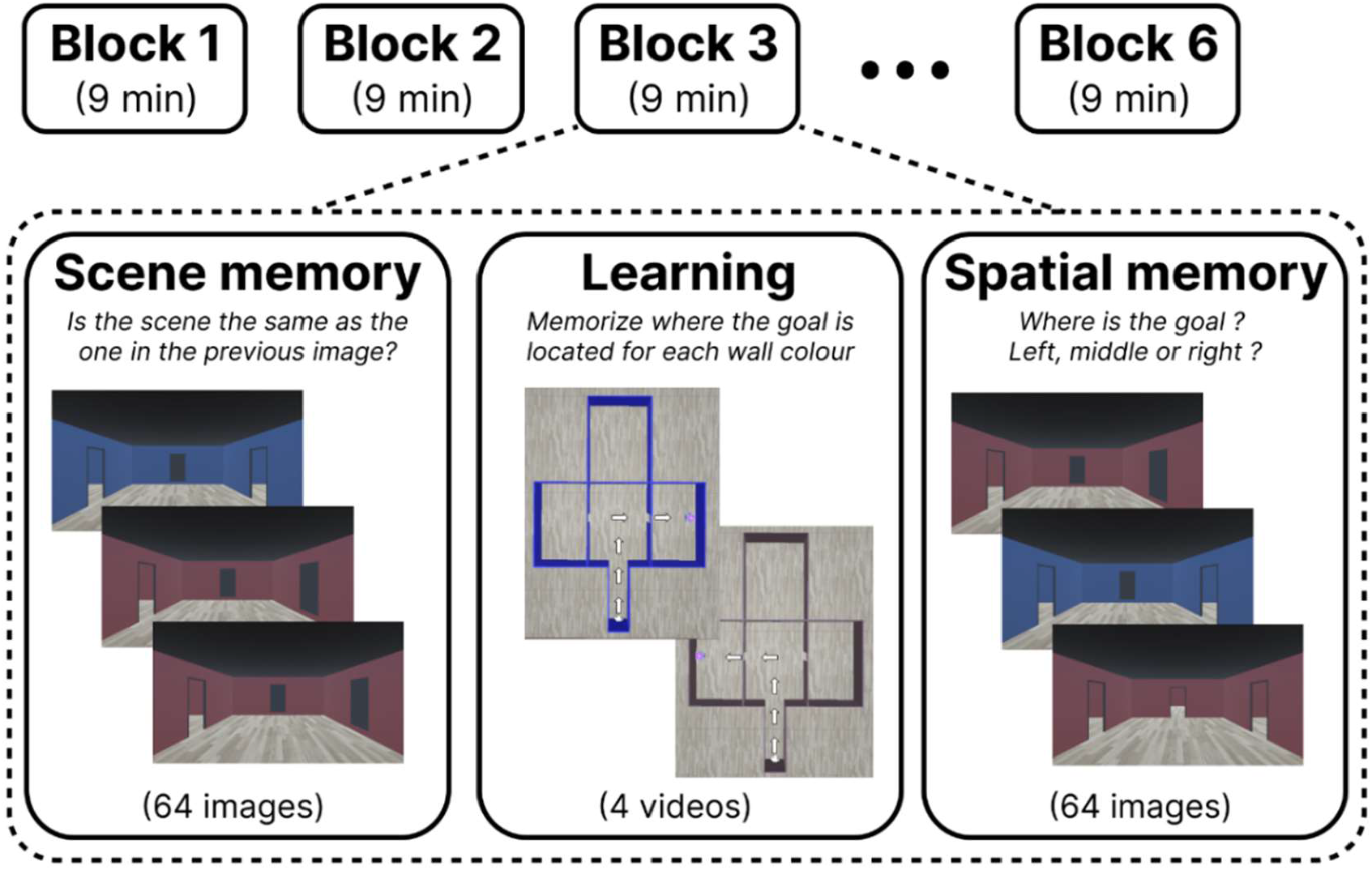
Presentation of the paradigm design and a subset of the stimuli used for the scene memory and spatial memory tasks. It is important to note that the images in the learning phase are used here only to illustrate the path followed during passive navigation, and that participants did not have access to this aerial view of the environment.

In both the scene memory and the spatial memory tasks, images were displayed for a fixed duration of 1 second, followed by a randomly jittered interstimulus fixation cross lasting between 0.5 and 1 second. During this interval, participants were provided with auditory feedback about whether they had given the right answer. Overall, each participant was presented with 1536 images (768 for each task), resulting in an average experimental duration of 55 minutes.

### EEG acquisition and processing

EEG signals were acquired at a sampling rate of 500 Hz using a 64-channel Waveguard™ original cap equipped with Ag/AgCl electrodes. The electrodes were connected to an eego™ mylab amplifier from ANT Neuro and digitized at a 24-bit resolution. The reference electrode was CPz, and the ground electrode was AFz. Impedances were kept below 20 kΩ, with most electrodes maintained below 10 kΩ. EEG recordings and stimulus presentations were synchronized using the LabStreamingLayer software Labrecorder (LSL Labrecorder version 1.13). EEG data was preprocessed offline using MATLAB (R2021a; The MathWorks Inc., Natick, MA, USA) with custom scripts from the EEGLAB toolbox version 14.1.0b (Delorme & Makeig, 2004). These scripts were modified from a previously established processing pipeline (Delaux et al., 2021) based on the BeMobil pipeline (Klug et al., 2022).

The data were first downsampled to 250Hz and the time points were corrected using a hardware trigger. Line noise was automatically removed using the recently developed Zapline-Plus algorithm (Klug & Kloosterman, 2022). Noisy channels were identified and rejected using the default parameters suggested in the PREP pipeline (Bigdely-Shamlo et al., 2015). On average, 2.93 channels (SEM = 0.33) were rejected per participant. The channels were reconstructed through spherical interpolation of nearby channels and subsequently referred to a common average. Artifact signals were automatically removed using Artifact Subspace Reconstruction (ASR) (Kothe & Jung, 2015), which used clear sections of the data to determine component rejection thresholds. An ASR cutoff parameter of 20 was used in line with the suggested optimal range of 10 to 100 (Chang et al., 2020). The cleaned dataset was temporally high-pass filtered at 1.5 Hz using a Hamming window Finite Impulse Response (FIR) filter with a transition bandwidth of 0.5 Hz, a passband edge of 1.25 Hz, and an order of 1650 (Klug & Gramann, 2021). Subsequently, the Adaptive Mixture Independent Component Analysis (AMICA) algorithm (Palmer et al., 2008) was applied to the dataset. Next, each independent component (IC) was assigned a dipolar source that was reconstructed using the equivalent dipole model (DipFit; Oostenveld & Oostendorp, 2002)). To classify components into 7 classes, we used the ICLabel algorithm (Pion-Tonachini et al., 2019) and chose default percentages for classification. We then removed heart, line noise, eye, and channel noise components. On average, we excluded 14.33 components (SEM = 0.82) per participant, similar to previous studies such as Harel *et al*. (2022) who excluded on average 14 components. Next, we applied a bandpass filter to the data, using a lower cutoff frequency of 0.3 Hz (with a transition bandwidth of 0.5 Hz, passband edge at 0.55 Hz, and an order of 1650) and an upper cutoff frequency of 80 Hz (with a transition bandwidth of 20 Hz, passband edge at 80 Hz, and an order of 42).

### Behavioral and EEG analyses

We extracted the reaction time (*i*.*e*., the time between image appearance and the answer of participants) and accuracy scores (*i*.*e*., the percentage of correct retrieval) for both scene and spatial memory tasks, and for the three different affordance conditions. We then applied linear mixed models to these extracted values using subjects as a random intercept.

When analyzing EEG recordings, we focused on posterior lateral electrodes associated with scene-selective regions used in previous studies (Hansen et al., 2018; Harel et al., 2016, 2020, 2022; Kaiser et al., 2020) corresponding to P5-P7-PO7 for the left hemisphere and P6-P8-PO8 for the right hemisphere. We extracted epochs and excluded the one with less than 90% artifact-free data, resulting in an average retention rate of 73.8% (mean number of retained epochs per participant: 1170; SEM = 36.73). To further enhance data quality, we used the *bemobil_reject_epochs* function (Klug et al., 2022), which eliminated 5% of the worst epochs for each participant, based on the sum of four measures (mean of channel means, mean of channel SDs, SDs of channel means, and SD of channel SDs). Subsequently, the epochs were baseline-corrected using the pre-stimulus interval from -200 milliseconds to 0 milliseconds (image presentation). Next, we identified the peak amplitude windows by extracting the latency of the peak for the grand average across all experimental conditions. These windows, with -50 milliseconds and +50 milliseconds padding, were applied to determine individual peak amplitudes and latencies for the P1, N1, and P2 components (peaking respectively on average at 125ms [SEM = 1.9]; 175ms [SEM = 2.2] and 242ms [SEM = 2.3]). We chose to use the mean peak amplitude, which is more robust to artefacts and corresponds to the average of the most positive value surrounded by two lower values (Luck, 2014).

Statistical analyses of EEG results were performed using R statistical software (version 4.2.1, R Foundation for Statistical Computing, Vienna, Austria) with R studio (version 2022.07.02+576) and linear mixed-effects models from the lme4 package (Bates et al., 2014). After comparing different model structures using the Akaike information criterion (AIC) (Akaike, 1974), we included task (scene memory *vs*. spatial memory), affordance condition (1, 2 and 3 doors) and hemisphere (left *vs*. right) as independent factors, with subjects as random intercept. To extract *p*-values, we used the *anova* function and the Satterthwaite method using a type III sum of square. Estimated marginal means (EMMs) were calculated using the *emmeans* package in R and correspond to the value reported hereafter. Finally, post-hoc Tukey’s honest significant difference (HSD) tests were performed, and partial eta squared, and Cohen’s d were reported as post-hoc effect size measures. To verify normality of the residuals and homoscedasticity, both were carefully inspected using quantile-quantile plots and box plots, respectively.

## Results

### Behavioral analysis

The reaction time results (**Figure 2.A**) revealed a statistically significant main effect of the task (F_(1,125)_ = 25.51, *p* < 0.001, η _p_ ^2^ = 0.15, 95% CI = [0.06, 0.26]). As predicted, participants responded faster to the scene memory task (Estimated Marginal Mean = 535 milliseconds, SE = 8.18 ms) than to the spatial memory task (EMM = 551 ms, SE = 8.18 ms). The number of doors presented had no significant effect on reaction time (F_(1,125)_ = 0.27, *p* = 0.76). To further investigate the potential effect of door position (*i*.*e*., left, center, and right) on behavior, we conducted a complementary analysis comparing reaction times when participants were presented with a single open door. This analysis revealed a significant main effect of door position on reaction time in both the spatial and scene memory tasks (F_(2, 125)_ = 6.03, p = 0.003, η _p_ ^2^ = 0.09, 95% CI = [0.01, 0.19]), with shorter reaction times observed when the open door was in the central position (EMM = 527ms, SE = 5.52ms) than in the left or right positions (EMM = 536ms, SE = 5.52ms and EMM = 538ms, SE = 5.52ms, respectively). In a complementary analysis, we used an ANOVA to investigate a possible effect of the deroulment of the experiment (F_(1,29)_ = 11.33, *p* = 0.002) and found a shorter reaction time for the last experimental block (M = 537ms, SE = 7.30) compared to the first block (M = 557ms, SE = 6.54), with a similar pattern for both tasks (F_(1,29)_ = 0.176, *p* = 0.678). These results suggests that participants improved their reaction time over the course of the experiment, highlighting a training effect.

**Figure 2.**
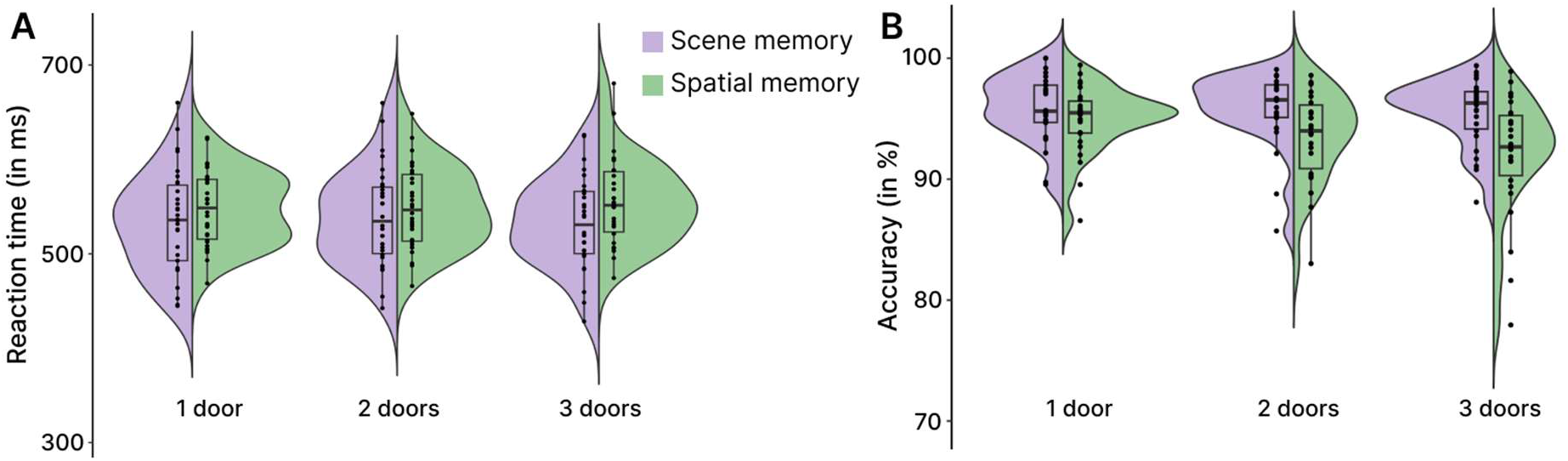
Split-violin plots of the behavioral data for scene memory tasks (purple) and spatial memory tasks (green) depending of the number of affordances available **A**. Plot of the reaction time after image presentation. **B**. Plot of the accuracy, corresponding to the percentage of correct participant answers.

The accuracy results (**Figure 2.B**) also showed that the task had a significant main effect (F_(1,125)_ = 41.82, *p* < 0.001, η _p_ ^2^ = 0.25, 95% CI = [0.13, 0.37]), with a higher accuracy during the scene memory tasks (EMM = 95.7%; SE = 0.58) than during the spatial memory tasks (EMM = 93.47%; SE = 0.58). The post-hoc analysis of the interaction between the number of open doors and the task (F_(2,125)_ = 4.52, *p* = 0.013, η_p_ ^2^ = 0.07, 95% CI = [0.00, 0.16]) revealed that the task related effect was only present in the 2-door condition (t_(125)_ = 3.80, *p* = 0.003, d = 1.05, 95% CI = [0.48, 1.62]) and 3-door condition (t_(125)_ = 5.83, *p* < 0.001, d = 1.62, 95% CI = [1.03, 2.20]), but not in the 1-door condition (t_(125)_ = 1.58, *p* = 0.615). These high performances are consistent with the relative simplicity of the paradigm and confirm that participants had acquired knowledge of the goal position during the spatial memory task. The number of open doors had a significant main effect (F_(2,125)_ = 7.55, *p* < 0.001, η ^2^ = 0.11, 95% CI = [0.02, 0.21]), but the effect was limited to the spatial memory task (t_(125)_ = 4.87, *p* < 0.001, d = 1.35, 95% CI = [0.77, 1.93]), and no effect was found for the scene memory task (t_(125)_ = 0.62, *p* = 0.99). The accuracy score was lower for the 3-door condition compared to the 1-door condition (t_(125)_ = 3.88, *p* < 0.001, d = 0.76, 95% CI = [0.35, 1.17]), but there were no significant differences between the 1-door and 2-door conditions (t_(125)_ = 1.81, *p* = 0.169), or between the 2-door and 3-door conditions (t_(125)_ = 2.07, *p* = 0.100).

In summary, the number of affordances modulated the behavior of the participants differently in the two tasks. While an increase in the number of affordances did not affect scene memory performance, it affected performance in the spatial memory task, which worsened as the number of affordances increased. This result suggests that the processing of affordances may have interfered with the spatial memory task, in which participants had more contextual information and were tasked to retrieve a goal behind one of the doors.

### EEG results of the main part of the experiment

#### Effect of the task, comparing scene memory and spatial memory for both hemispheres

In this first analysis, we compared the grand average of ERPs generated by the scene memory and spatial memory tasks (**Figure 3.A**) with the extracted mean peak values (**Figure 3.B**).

**Figure 3.**
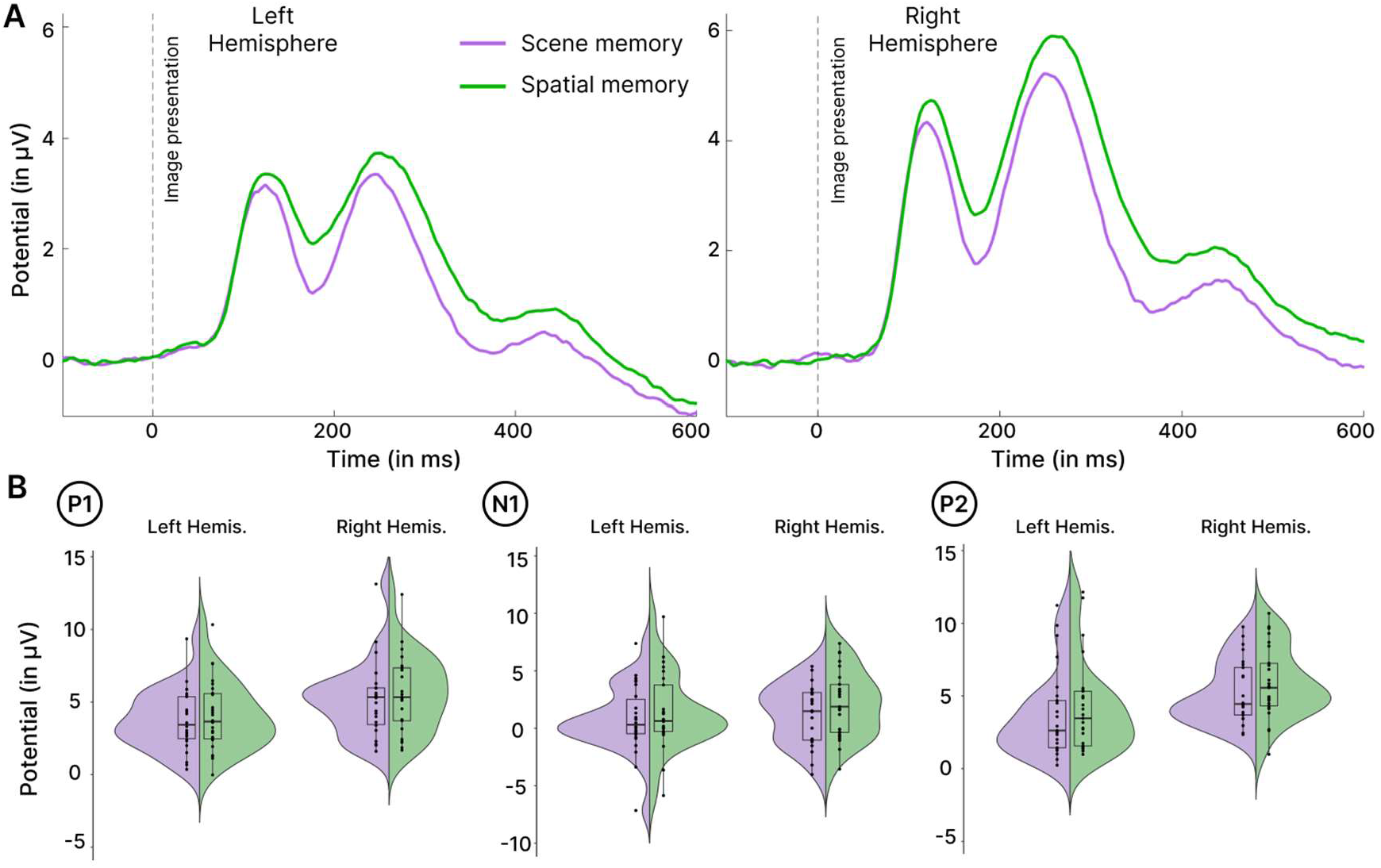
ERP results for scene memory tasks (purple) and spatial memory tasks (green) **A**. Grand-average ERPs measured over the P6-P8-PO8 electrodes on the right hemisphere and over the P5-P7-PO7 electrodes on the left hemisphere **B**. Split violin plots of the extracted individual amplitudes for the P1, N1, and P2 components in scene memory tasks (purple) and spatial memory tasks (green).

The number of epochs extracted was similar for both tasks (t_(25)_ = 1.30, *p* = 0.205). As predicted, the P2 amplitude was higher in the spatial memory task than in the scene memory task (F_(1,73.15)_ = 4.33, *p* = 0.04, η_p_ ^2^ = 0.06, 95% CI = [0, 0.18]), suggesting a modulation of prior spatial knowledge on the early markers of scene processing. On the contrary, the N1 amplitude was more negative in the scene memory task than in the spatial memory task (F_(1,73.54)_ = 6.45, *p* = 0.013, η_p_ ^2^ = 0.08, 95% CI = [0, 0.22]). Finally, the P1 amplitude did not differ between the tasks (F_(1,74.05)_ = 2.07, *p* = 0.15). Considering latencies, we reported a delayed P2 component in the spatial memory task (F_(1,273.97)_ = 22.51, *p* < 0.001, η_p_ ^2^ = 0.08, 95% CI = [0.03, 0.14]), while there was no difference for the P1 and N1 components (F_(1,274.02)_ = 1.12, *p* = 0.291 and F_(1,274.01)_ = 2.57, *p* = 0.110, respectively). We also reported lateralization differences for the P1 and P2 components that were higher in the right hemisphere compared to the left hemisphere (P2 component: F_(1,73.14)_ = 47.49, *p* < 0.001, η_p_ ^2^ = 0.39, 95% CI = [0.23, 0.53]; P1 component: F_(1,74.05)_ = 34.71, *p* < 0.001, η_p_ ^2^ = 0.32, 95% CI = [0.16, 0.47]). No hemisphere effect was reported for the N1 component (F_(1,73.07)_ = 3.14, *p* = 0.080). This overall greater activity in the right hemisphere is consistent with a previous ERP study on navigational affordance processing during passive scene perception (Harel et al., 2022) and a previous fMRI study on OPA activity during visually guided navigation (Persichetti & Dilks, 2018).

#### Effect of the number of doors

This analysis focused on the comparison between the grand-average ERPs based on the number of doors and investigated the possible modulation of the ERP component by the number of navigational affordances (1-, 2-, and 3-door condition) in both the scene memory (**Figure 4.A**) and spatial memory tasks (**Figure 4.B**).

**Figure 4.**
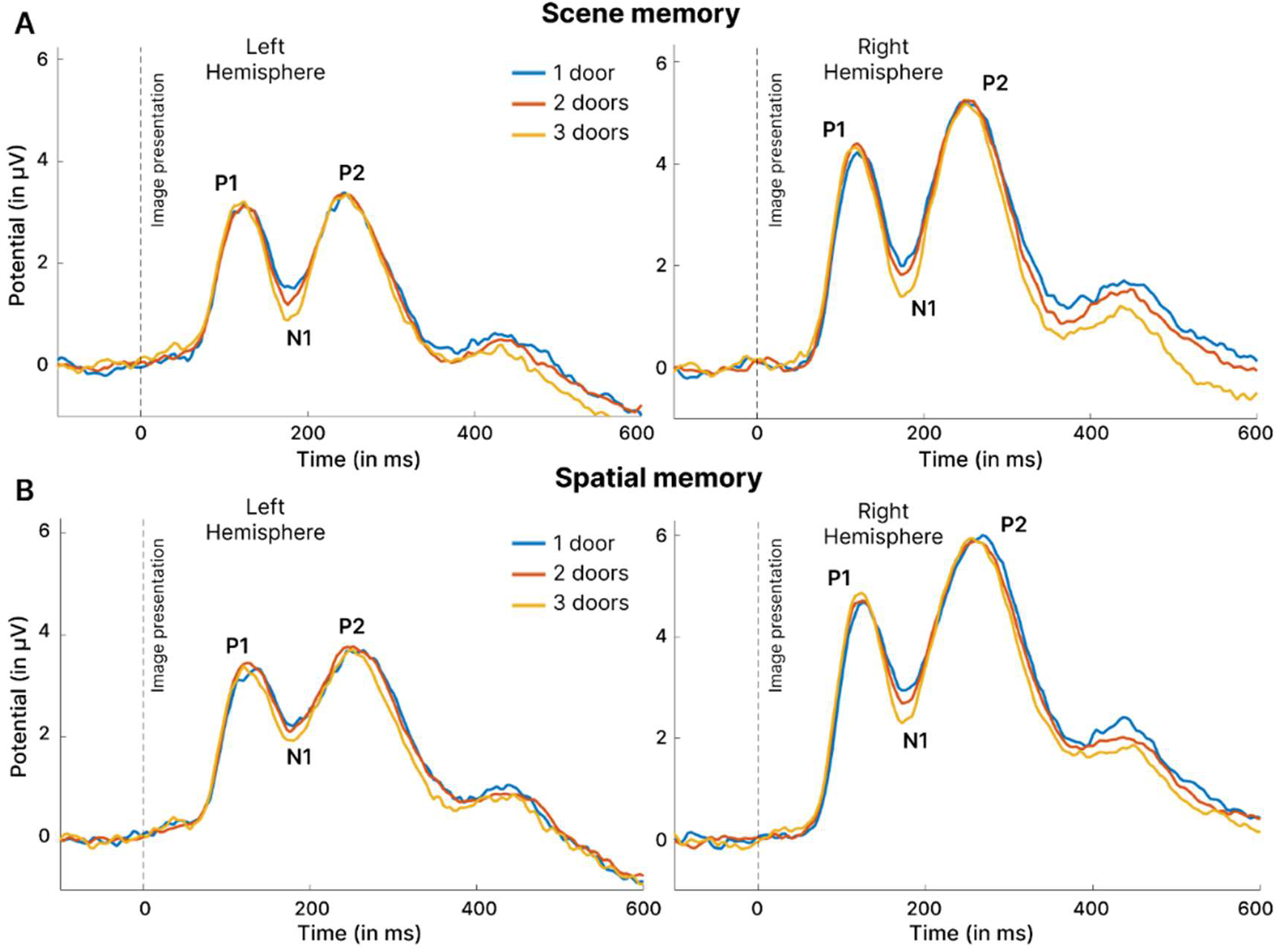
ERP results comparing the activity in the 1-door, 2-door and 3-door condition between the left and right hemisphere. Statistics were computed using the extracted peaks amplitude and latencies. **A**. In the scene memory task. **B**. In the spatial memory task.

Our first results showed no interaction between the affordance conditions and the task, suggesting that both tasks generated a similar pattern of results in the different affordance conditions (P1: F_(2,275)_ = 0.057, *p* = 0.99; N1: F_(2,274.01)_ = 0.0032, *p* = 0.99; P2: F_(2,274)_ = 0.017, *p* = 0.98). Contrary to expectations and previous results (Harel et al., 2022), we reported no modulation of the P2 component by the number of affordances in both the scene and spatial memory tasks (F_(2,274)_ = 0.005, *p* = 0.99). The P1 amplitude was also not modulated by the number of affordances (F_(2,275)_ = 0.159, *p* = 0.85), while the N1 showed a main effect of the number of affordances (F_(2,274.01)_ = 3.62, *p* = 0.028, η_p_ ^2^ = 0.03, 95% CI = [0, 0.07]), with the 3-door condition associated with a more negative amplitude than the 1-door condition only (t_(274)_ = 2.63, *p* = 0.024, d = 0.37, 95% CI = [0.08, 0.65]). According to the literature, this N1 modulation is intriguing and could be due to a difference in the orientation of attention related to the position of the doors (Hillyard et al., 1998).

To assess the potential influence of door location on the previously observed results and notably the modulation of the N1 component by the number of affordances, we analyzed the interaction between task and door location. In **Figure 5**, we pooled the spatial and scene memory ERP representations for clarity, as no interaction had been reported between task and door location for P1 (F_(2,124.02)_ = 0.91, *p* = 0.403), N1 (F_(2,124.02)_ = 0.51, *p* = 0.604) or P2 (F_(2,125)_ = 1.86, *p* = 0.160), which indicates a similar pattern of activity for both tasks. The following results were computed using the extracted peaks from **Figure 5.A**.

**Figure 5.**
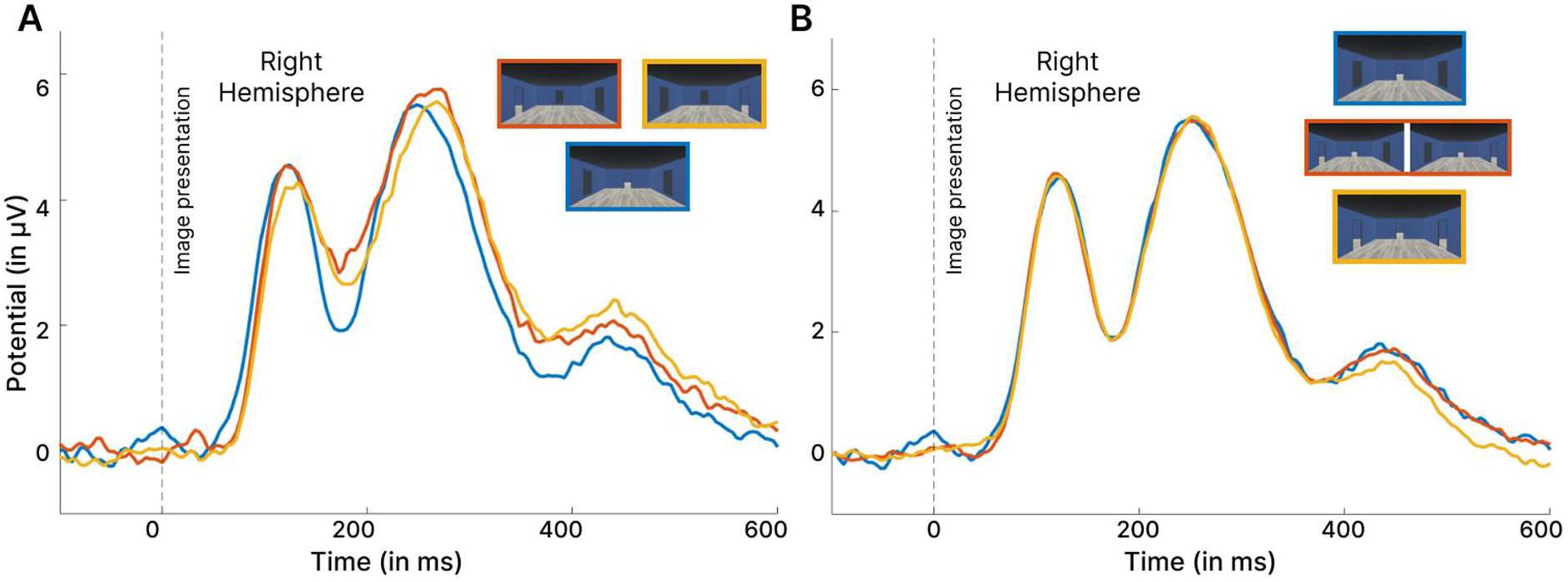
ERP results for the right hemisphere merging scene memory and spatial memory tasks. **A**. ERP comparison in the 1-door condition, with either the left, right or central door open. Statistical analyses were conducted with the peaks extracted from this comparison. **B**. Complementary analysis representing the ERP comparison in all the door conditions in which the central door is open.

The P1 and P2 amplitudes did not show any effect of door location (F_(2,124.02)_ = 0.82, *p* = 0.443 and F_(2,125)_ = 1.57, *p* = 0.212, respectively), but the N1 amplitude reflected a significant main effect of door location (F_(2,124.01)_ = 9.96, *p* < 0.001, η_p_ ^2^ = 0.14, 95% CI = [0.04, 0.25]). The post-hoc analyses revealed that the N1 amplitude was more negative with the central door than with the left or right door (t_(124)_ = 4.203, *p* < 0.001, d = 0.83, 95% CI = [0.41, 1.25] and t_(124)_ = 3.393, *p* = 0.002, d = 0.67, 95% CI = [0.26, 1.07], respectively), but there was no difference between the left and right door (t_(124)_ = 0.830, *p* = 0.685). These results support the idea that the modulation of the N1 component was not related to the number of affordances, but rather to the location of the door, with the open central door driving this effect. However, the absence of modulation of the P2 component remained intriguing and led to the follow-up analysis.

### ERP results of a supplementary control experiment

The lack of P2 modulation was surprising in light of the previous ERP study on early affordance processing by Harel *et al*. (2022). This discrepancy with our results could be influenced by the fact that participants in both our scene and spatial memory tasks were engaged in active scene processing (*i*.*e*., scene identification using wall color) as opposed to the passive scene perception in the previous work. To address this possibility, we conducted an additional ERP experiment with 20 new young participants, 17 remaining after the exclusion of 3 due to artifacted data (mean age: 23.76 years; SEM = 0.40; range: 21-27; 9 female), who performed a passive perception task. The same rooms (1-door, 2-door, and 3-door) were used as stimuli with the addition of a red fixation cross centered in the image, following the methodology of Harel *et al*. (2016, 2022). Participants were instructed to use the keyboard to indicate whether the horizontal or vertical bar of the cross had increased by 25%. The same number of images (1536, divided into 6 blocks) were presented in this passive perceptual task, with presentation times similar to those used in the scene memory and spatial memory tasks. We then compared the ERPs generated by a passive scene perception task in the 1-, 2-, and 3-door conditions (**Figure 6**).

**Figure 6.**
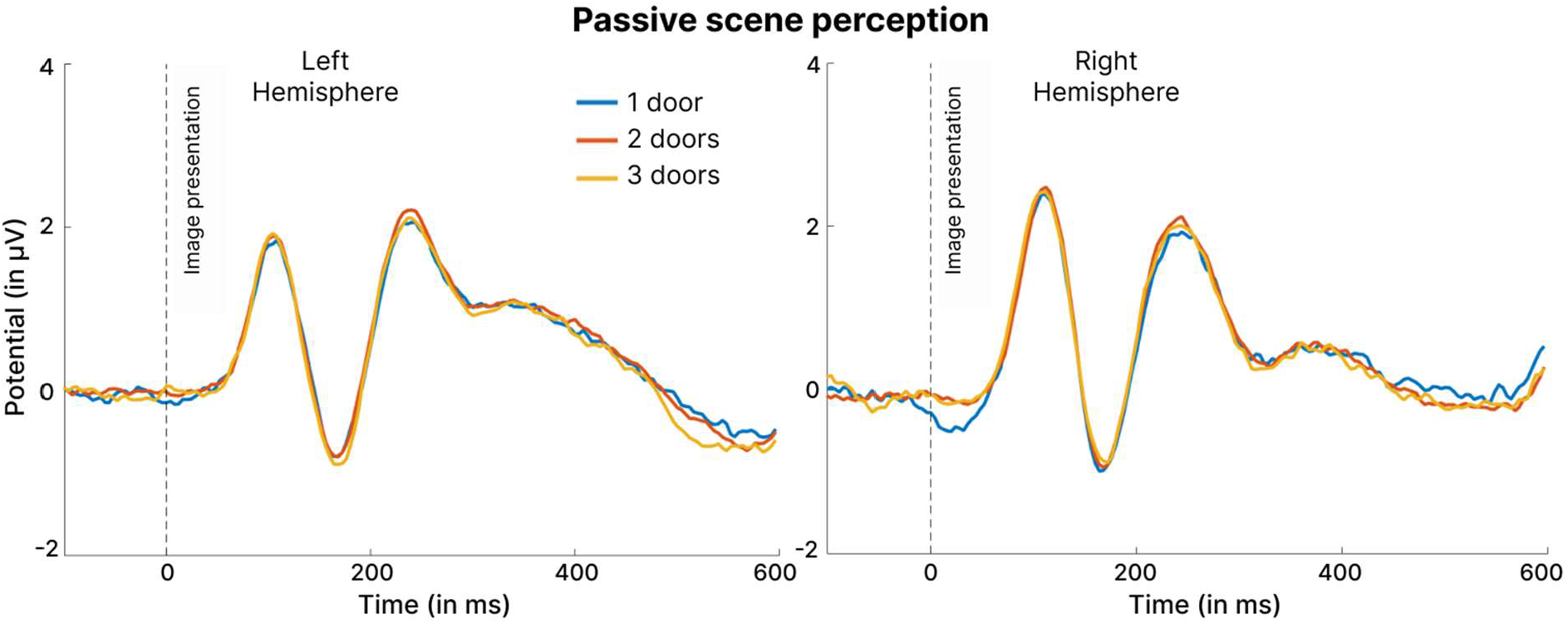
ERP results for the left and right hemispheres of our additional experiment involving a passive scene perception task depending on the number of open doors available. Statistical analyses were conducted on the peaks extracted from these results.

Linear mixed model analysis of the extracted mean peak amplitudes revealed that the number of open doors did not significantly affect the amplitude of any ERP component (all *p* > 0.59). Furthermore, there was no interaction between doors and hemisphere for any amplitude (all *p* > 0.47). Consistent with the previous part of the experiment, these results suggest that, in our experimental protocol, the number of affordances does not appear to affect the P1 or N1 components, and, more surprisingly, does not affect the P2 component.

## Discussion

In the current study, we aimed to investigate whether prior spatial knowledge of the immediate visual environment affects behavior and the associated early neural dynamics, while also investigating the interaction of this knowledge with the processing of navigational affordances. Using a univariate ERP analysis, we found that the amplitude of the scene-selective P2 component (Hansen et al., 2018; Harel et al., 2016, 2020, 2022; Kaiser et al., 2020) was higher in a spatial memory task in which participants had more contextual knowledge of the scene than in a scene memory task. Moreover, contrary to our initial hypothesis regarding the previous study by Harel *et al*. (2022), we did not report any effect of the number of visible exits on the scaling of P2 peak amplitude in either task, even in a supplementary experiment in which we used a passive scene perception task similar to theirs. Despite this absence of difference in the amplitude of ERPs with respect to the number of navigation possibilities, our behavioral results revealed a significant decrease in participants’ accuracy rate when the number of potential paths increased, but only in the spatial memory task. Taken together, our results emphasize an intricate influence of top-down processing related to prior spatial knowledge about the environment, on the early stages of visual processing, as well as bottom-up processing of the potential for navigation in a scene, highlighting the complex interactions between perceptual and spatial memory systems (Martin & Barense, 2023).

Our finding that the amplitude of the P2 component increased and was delayed when participants had to retrieve the goal position compared to a scene memory task in the same environment, supports the hypothesis that previously learned contextual information modulated even the earlier visual processing. Our results seem to be in line with previous work suggesting that the occipito-parietal P2 component plays a key role in mediating the comparison of visual input with information stored in memory (Cepeda-Freyre et al., 2020; Freunberger et al., 2007; Lefebvre et al., 2005). This was recently reinforced by Enge *et al*. (2023), who showed that the availability of prior semantic knowledge about an object’s function can immediately influence early cortical markers of that object’s processing, including the higher-level visual perceptual processing that occurs around 200 ms (Greene & Hansen, 2020; Groen et al., 2018). In a similar way, Klink *et al*. (2023) proposed that familiarity with a scene was reflected in a peak of EEG activity during the early stages of visual processing (after 250 ms) in the same time range as the P2 component, suggesting an early deeper processing of already learned information compared to novel stimuli. These results, interpreted as a potential top-down influence, are consistent with recent fMRI studies showing the presence of a functional brain network composed of “place-memory areas” immediately adjacent to the SSRs, which are proposed to integrate the visuospatial context of a scene and the perceived visual stimulus (Steel et al., 2021, 2023, 2024). These converging findings suggest that memory-derived information may play a more important role than the conventional view of scene processing suggests. In this sense, our ERP results also seem to support this idea, with a possible increased activity over the region of the scene-selective network when participants have more spatial contextual knowledge about the currently perceived scene.

Moreover, these task-related differences in brain activity are also associated with behavioral results showing that participants took slightly longer to respond in the spatial memory task. These results may indicate a differential monitoring of cognitive resources allocated to the identification task when remembered visuospatial information from the scene is added, as suggested by the reduced N1 amplitude in this task (Hillyard et al., 1998; Hopf et al., 2002; Warbrick et al., 2014; Wiegand et al., 2014). However, this N1 modulation may also reflect a reduced need for perceptual analysis, possibly as a result of prior training (Song et al., 2007; Y. Wang et al., 2010), due to our experimental design in which the spatial memory task was always performed after the scene memory task. Also, the longer reaction time reported here seems to reflect the involvement of additional cognitive processes related to differences between tasks in the level of hierarchical encoding of memory information (Hommel, 2000; McNamara et al., 1989; Peer & Epstein, 2021; Wiener & Mallot, 2003). Indeed, we could consider that the need to remember the position of a goal in the scene implies an additional hierarchical sublevel compared to the 1-back task performed in the scene-memory condition. In this sense, in the latter case, the participant can remain at the broader contextual level of the scene given by the color of the walls without having to access any encapsulated knowledge. Another possible explanation could lie in differential preattentional demands associated with the task or with the fixed order of presentation in our protocol, in which the spatial memory task is always presented after the scene memory task. However, this interpretation can be ruled out considering the similar amplitude of the P1 component between tasks. This component has been proposed as an early indicator of attentional control that reflects visuospatial attentional processing (Di Russo et al., 2003; Fu et al., 2010) and allows for efficient early stimulus categorization (Freunberger et al., 2007; Klimesch, 2011). P1 amplitude results thus suggest that both tasks involve comparable attentional demands and indicate comparable engagement of participants in both tasks.

A second aim of the current study was to investigate the role of prior spatial knowledge on the automatic processing of navigational affordances, as reflected by modulation of the P2 amplitude, as proposed in a previous ERP study (Harel et al., 2022). As hypothesized, our ERP results showed that the P2 component was not modulated by the number of visible paths in the spatial memory task, suggesting that prior knowledge about the goal to be reached could influence the automatic extraction of navigational affordances. However, in contrast to Harel *et al*. (2022), we also reported a lack of modulation of the P2 component during the scene memory task. A possible explanation for this surprising finding could be that performing a detection task in the visual scene could interfere with the automatic processing of affordances and mask the modulation of the P2 component by the number of affordances. This interference has been proposed in the context of visual object processing, where the participant’s behavioral goal influences the representation of visual objects in a top-down manner (Harel et al., 2014), and this could then explain our lack of P2 modulation. In the present study, we equated our two tasks in terms of perceptual load demand (*i*.*e*., a wall color discrimination task). To control for a possible task-related effect, we then reused our stimuli in a supplementary control experiment, but this time with the same passive perceptual task as proposed by Harel *et al*. (2022). We still reported no modulation of the P2 component by the number of affordances. This discrepancy is particularly noteworthy given the similarities in our experimental paradigms and stimuli. However, our findings appear to be consistent with previous univariate fMRI results that suggested no differences in OPA activity when the authors varied the number of available pathways while controlling for scene complexity by replacing missing doors with paintings (Bonner and Epstein., 2017). The authors interpreted this finding as a reflection of the visual complexity affecting the linear trend of activity in these regions, in line with preliminary work suggesting that the OPA is associated with local elements of scenes (Kamps et al., 2016). This difference in stimulus complexity between our task and the study by Harel *et al*. (2022) could then potentially explain the divergence in our results. This interpretation was also suggested by Dwivedi *et al*. (2024), who reported that the processing of navigational affordances occurred later, and they explained this difference in latency by the nature of their stimulus or the task being performed.

Despite an absence of modulation of the early P2 amplitude in posterior parietal electrodes, analysis of the behavioral data revealed an interesting pattern associated with the impact on performance of the number of affordances. Contrary to our expectations, increasing the number of available doors specifically decreased accuracy in the spatial memory task, while scene memory was unaffected. This result suggests that the participants had to consider other action possibilities offered by the environment even though they already had a good knowledge of the goal position, as shown by their very high accuracy rate. In other words, this behavioral pattern suggests an interaction between prior spatial knowledge about the environment and the automatic processing of the available navigational affordances (Bonner & Epstein, 2017; Harel et al., 2022; Patai & Spiers, 2021). Future work using eye-tracking could help to explore this interaction in greater detail, by investigating, for example, whether participants looked at the exits differently depending on the task they were performing (Henderson & Hollingworth, 1998; Mills et al., 2011). Taken together, our results challenge the notion that the P2 component is only influenced by the number of navigational affordances available and highlight the complexity of this component and its role in visual scene processing (Freunberger et al., 2007; Hansen et al., 2018; Harel et al., 2020; Mecklinger et al., 2009; Philips & Takeda, 2009). Our behavioral results also highlight the intricate influence of prior spatial knowledge and the proposed automatic processing of navigational affordances, but this was not reflected in early visual brain activity. This pattern of results aligns with what was reported by Wang *et al*. (2023), who also investigated the differences in the processing of navigational affordances when participants performed different tasks, whether virtually navigating through the environment or not. They are in line with the previous findings of Djebbara *et al*. (2019) suggesting that in the later sensorimotor stages, the processing of affordances may be context dependent rather than automated, with brain dynamics modulated by the width of affordances only when participants know they have to interact with the environment. They also suggested that the perception of affordances was divided into early stimulus-driven and later goal-driven processes, forming a continuum (Djebbara et al., 2022). Despite differences between their protocol design and ours, our behavioral results also seem to point in this direction, with a possible specific goal-directed processing of the affordances that occurred only when participants were tasked with reaching a goal hidden behind an open door, which affected their performance when they were offered more motor options.

Finally, we reported a shorter reaction time when the central door was open in both of our tasks. This behavioral result was associated with an increased amplitude of the N1 component. In our experiment, the central door corresponded to the location on which attention was focused when the fixation cross was presented during the inter-trial interval. Thus, the increased N1 amplitude when the central door was open is consistent with the notion that directing attention to a specific location enhances the N1 response elicited by objects presented at attended locations (Hillyard & Anllo-Vento, 1998; Vogel & Luck, 2000). Interestingly, no difference was observed when the door was displayed on the left or right side, suggesting that the open doors were not subject to directed visual processing. If directed visual processing had occurred, we would have observed a contralateral N1, with increased N1 amplitude in the right hemisphere in the left door condition, as has been extensively reported in ERP research (Wascher et al., 2009). Consequently, we propose that the increased N1 in the central door condition was not due to the task itself, but to the fact that participants were able to perform the discrimination task using the central wall when the central door was open, allowing for more efficient processing in the light of attention. This interpretation is further supported by the lack of modulation of the N1 component observed in our control experiment when participants had to perform a fixation cross task.

Although this represents an interesting pattern of results, this N1 modulation may have affected the results we obtained for the P2 component, and future work could consider controlling for this limitation, by placing the doorways in different parts of the scene, for example, and avoiding the central location. Such work could also consider the use of source reconstruction to gain insight into the brain regions involved. This would be interesting because despite the selection of electrodes associated with scene-selective regions based on previous work (Harel et al., 2016, 2022; Kaiser et al., 2020), the spatial resolution of EEG does not allow us to draw definitive conclusions about whether the reported activity originates from the OPA, PPA, or more anterior regions, as reported by Steel *et al*. (2021). However, recent methodological advances, particularly the incorporation of fMRI-derived priors (Abreu et al., 2022; Cottereau et al., 2015) appear to offer the possibility of improving spatial accuracy (Liu et al., 2023). This may allow future work to distinguish between OPA, PPA, and MPA activity, as was recently achieved by Durteste *et al*. (2023), or to investigate the role of place-memory areas (Steel et al., 2021, 2023).

In summary, this work proposes that prior spatial knowledge about a scene, such as the location of a goal in the environment, modulates early cortical activity associated with scene-selective regions. This memory information may also interact with bottom-up processing of scene content, such as navigational affordances. This finding reinforces the idea that memory information may have a more profound influence on the overall function of early visual processing than previously suggested based on the conventional hierarchical visual processing perspective.

## CRediT author statement

**Naveilhan Clément:** Conceptualization, Methodology, Formal analysis, Investigation, Writing – Original Draft; **Sauley-Caret Maud:** Methodology, Investigation,; **Raphaël Zory:** Writing - Review & Editing, Funding acquisition; **Stephen Ramanoël:** Conceptualization, Supervision, Writing – Review & Editing, Funding Acquisition, Project administration.

## Acknowledgements

This research would not have been possible without the generous and dedicated participation of the volunteer subjects who generously donated their time and effort. The authors are grateful for their participation.

This work was supported by the French government through the UCA^JEDI^ project managed by the National Research Agency (ANR-15-IDEX-01) and, in particular, by the interdisciplinary Institute for Modeling in Neuroscience and Cognition (NeuroMod) of Université Côte d’Azur.

## Code and data availability

The complete set of raw data, codes, and stimuli generated for the analyses presented in this paper can be accessed through the OSF repository at: https://osf.io/5cqxy/?view_only=2957dac1a1304c2db6e1fb3b056c5008

## Gender Citation Balance Indices

The Gender Citation Bias Index (GCBI; Fulvio et al., 2021) values calculated indicated significant imbalances in citations: Male-Male (MM) pairings showed moderate overcitation with an index of 0.53, Male-Female (MF) showed undercitation with an index of -0.49, Female-Male (FM) pairings showed overcitation with an index of 0.65, and Female-Female (FF) pairings showed significant undercitation with an index of -0.83.

